# Network analysis reveals a distinct axis of macrophage activation in response to conflicting inflammatory cues

**DOI:** 10.1101/844464

**Authors:** Xiaji Liu, Jingyuan Zhang, Angela C. Zeigler, Anders R. Nelson, Merry L. Lindsey, Jeffrey J. Saucerman

**Affiliations:** Department of Biomedical Engineering, University of Virginia, Charlottesville, VA, USA; Department of Cellular and Integrative Physiology, University of Nebraska Medical Center and Research Service, Nebraska-Western Iowa Health Care System, Omaha, NE 68198, USA

## Abstract

Macrophages are subject to a wide range of cytokine and pathogen signals in vivo, which contribute to differential activation and modulation of inflammation. Understanding the response to multiple, often conflicting, cues that macrophages experience requires a network perspective. Here, we integrate data from literature curation and mRNA expression profiles to develop a large-scale computational model of the macrophage signaling network. In response to stimulation across all pairs of 9 cytokine inputs, the model predicted activation along the classic M1-M2 polarization axis but also a second axis of macrophage activation that distinguishes unstimulated macrophages from a mixed phenotype induced by conflicting cues. Along this second axis, combinations of conflicting stimuli, interleukin 4 (IL4) with lipopolysaccharide (LPS), interferon-γ (IFNγ), IFNβ, or tumor necrosis factor-α (TNFα), produced mutual inhibition of several signaling pathways, e.g. nuclear factor κB (NFκB) and signal transducer and activator of transcription 6 (STAT6), but also mutual activation of the phosphoinositide 3-kinases (PI3K) signaling module. In response to combined IFNγ and IL4, the model predicted genes whose expression was mutually inhibited, e.g. inducible nitric oxide synthase (iNOS) and arginase 1 (Arg1), or mutually enhanced, e.g. IL4 receptor-α (IL4Rα) and suppressor of cytokine signaling 1 (SOCS1), which was validated by independent experimental data. Knockdown simulations further predicted network mechanisms underlying functional crosstalk, such as mutual STAT3/STAT6-mediated enhancement of IL4Rα expression. In summary, the computational model predicts that network crosstalk mediates a broadened spectrum of macrophage activation in response to mixed pro- and anti-inflammatory cytokine cues, making it useful for modeling in vivo scenarios.

**Summary sentence:** Network modeling of macrophage activation predicts responses to combinations of cytokines along both the M1-M2 polarization axis and a second axis associated with a mixed macrophage activation phenotype.

## Introduction

Macrophages are central mediators of inflammation across a diverse range of protective or pathogenic processes including antimicrobial defense, anti-tumor immune responses, allergy and asthma, wound healing, and autoimmunity.[1]–[6] Tumor-associated macrophages generally exhibit an anti-inflammatory phenotype in response to hypoxic tumor microenvironment signals.[5] In rheumatoid arthritis, both pro- and anti-inflammatory cytokines stimulate macrophages to control inducible nitric oxide synthase (iNOS) expression and nitric oxide production, which is implicated in inflammation, angiogenesis, and tissue reconstruction.[6] After myocardial infarction, the macrophage population consists of subtypes that regulate the early pro-inflammatory and later anti-inflammatory reparative phases of infarct remodeling. Pro-inflammatory macrophages mediate the release of pro-inflammatory cytokines, whereas anti-inflammatory macrophages mainly participate in wound-healing.[7]–[11]

Macrophage infiltration into tissue and activation are coordinated by a variety of chemokines and cytokines. These environmental cues induce different macrophage phenotypes, characterized by distinct gene expression patterns and cell functions. Historically, macrophages in vitro have been classified into the classically (pro-inflammatory, M1) activated and the alternatively (anti-inflammatory, M2) activated phenotypes, each associated with specific markers. Lipopolysaccharide (LPS) and interferon-γ (IFNγ) are the prototypical stimuli for M1-type activation, and interleukin(IL) 4 is a prototypical M2-type stimulus.[12], [13] However, a number of studies have shown more diverse, stimulus-dependent macrophage phenotypes.[1], [4], [14]–[17] *In vivo* studies further indicate that macrophages respond to more complex, tissue-specific combinations of signaling factors than typically studied in vitro.[18], [19] Several recent reviews have noted that macrophage activation, orchestrated by complex spatiotemporally signaling cues, extends well beyond the linear M1/M2 spectrum and requires reassessment of current conceptual models.[20]–[22]

Developing more accurate conceptual models will require comprehensive assessments of macrophage phenotypes and systems biology frameworks that mechanistically link cues to phenotype. Advances in transcriptomics have provided genome-scale signatures of macrophage responses that extend beyond the limited marker panels previously considered. Omics studies have been critical in defining the complexity of macrophage responses that depend on cell source, timepoints of evaluation, and stimuli applied.[14], [23], [24] Network models are needed to mechanistically explain how complex cytokine inputs produce such signatures. [25], [21], [26] Large-scale network models have previously revealed key signaling properties of a number of mammalian cell types, including cardiac myocytes, fibroblasts, and T cells.[27]–[29]

To address this challenge, here we developed a large-scale, logic-based differential equation (LDE) computational model of macrophage activation. We refined and validated the model semi-quantitatively using RNA-Seq data from LPS+IFNγ or IL4-stimulated peritoneal macrophages. To examine how this network resolves conflicting cytokine cues, as often occurs in vivo, we simulated all pairwise combinations of 9 cytokine inputs. Predictions of gene expression in response to combined IFNγ and IL4 treatment were validated against an independent RNA-Seq dataset, and comprehensive knockdown simulations were used to identify underlying crosstalk mechanisms.

## Results

### Developing a large-scale, logic-based differential equation model of the macrophage activation signaling network

We performed a manual literature curation of the macrophage activation signaling network, integrating signaling pathways from review articles, original research articles, and a previous computational model (see **Methods**).[1], [30]–[32] This curated signaling network incorporated 9 cytokine inputs, including the classic M1-inducing LPS and IFNγ, M2-inducing IL4, as well as 7 other cytokines important in macrophage activation: IFNβ, IL1, IL6, IL10, IL12 and tumor necrosis factor-α (TNFα).[33], [13], [30], [15] A total of 39 mRNAs were selected as model outputs based on their association with macrophage polarization in previous studies and their differential expression in murine peritoneal macrophages stimulated by either LPS+IFNγ or IL4 for 4h.[34] Transcriptional feedback was incorporated for expression of IκBα, IL4Rα and autocrine cytokines IFNβ, IFNγ, IL1, IL6, IL10, IL12, and TNFα. Overall, this signaling network included 139 nodes (mRNA, proteins, and small molecules) connected by 200 reactions (**Figure 1**). Using this network structure, a logic-based differential equation (LDE) model of this signaling network was automatically generated as previously described (see **Methods**).[35]–[37] A full description of model structure, parameters, and supporting literature is provided in **Supplementary Table S1**.

**Figure 1.**
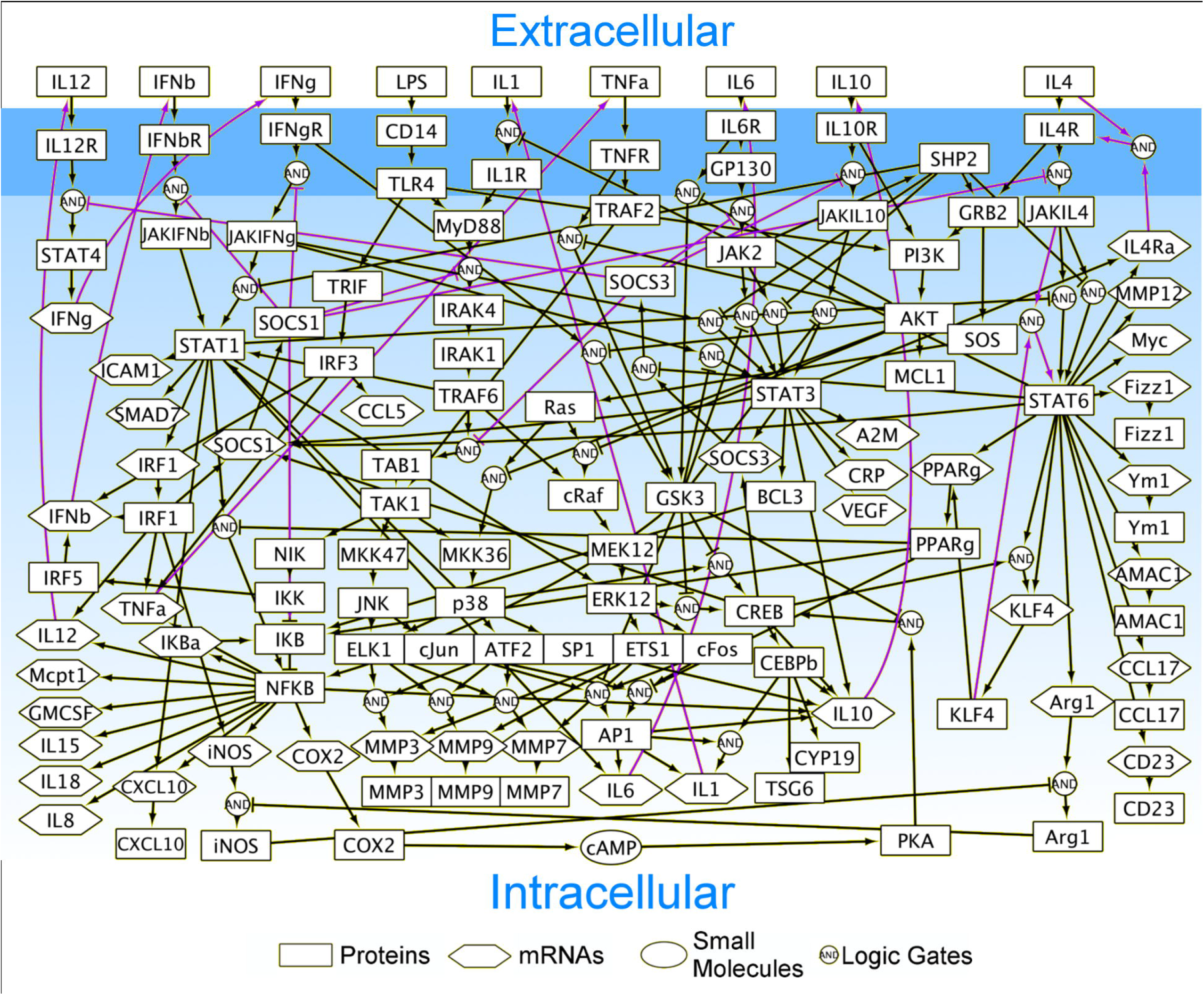
Network model of the peritoneal macrophage signaling network. Each of node represents a protein (rectangle), mRNA (hexagon), or small molecule (ellipse) in the network model. Each arrow indicates an activating (pointed arrow) or inhibiting (flathead arrow) reaction. Purple arrows highlight feedback reactions. Reactions involving multiple reactants were combined via AND logic gate (circled box). Multiple reactions affecting the same product were combined using OR gate logic. To simplify visualization, the translated nodes were overlapped under the corresponding signaling node (*e.g.* translated IL1 node covered by IL1 protein).

### Predicting signaling and gene expression dynamics in response to pro- and anti-inflammatory stimuli

The model was used to predict the dynamics of macrophage gene expression in response to stimulation by either pro-inflammatory LPS+IFNγ or anti-inflammatory IL4 (**Figure 2A**). Consistent with previous studies, genes used as pro-inflammatory phenotype markers such as IL1 and iNOS mRNAs were specifically induced by LPS+IFNγ stimulation, while anti-inflammatory markers such as arginase 1 (Arg1) mRNA were specifically induced by IL4 stimulation. IL1, IκBα, and matrix metallopeptidase 3/7/9 (MMP3/7/9) mRNAs were predicted to exhibit adaptive expression due to negative feedback regulation. Suppressor of cytokine signaling 1 (SOCS1) expression was predicted to increase under both conditions, but somewhat more strongly with LPS+IFNγ (**Figure 2B**). Network-wide responses to LPS+IFNγ and IL4 stimulation are visualized in **Supplementary Figure S1**.

**Figure 2.**
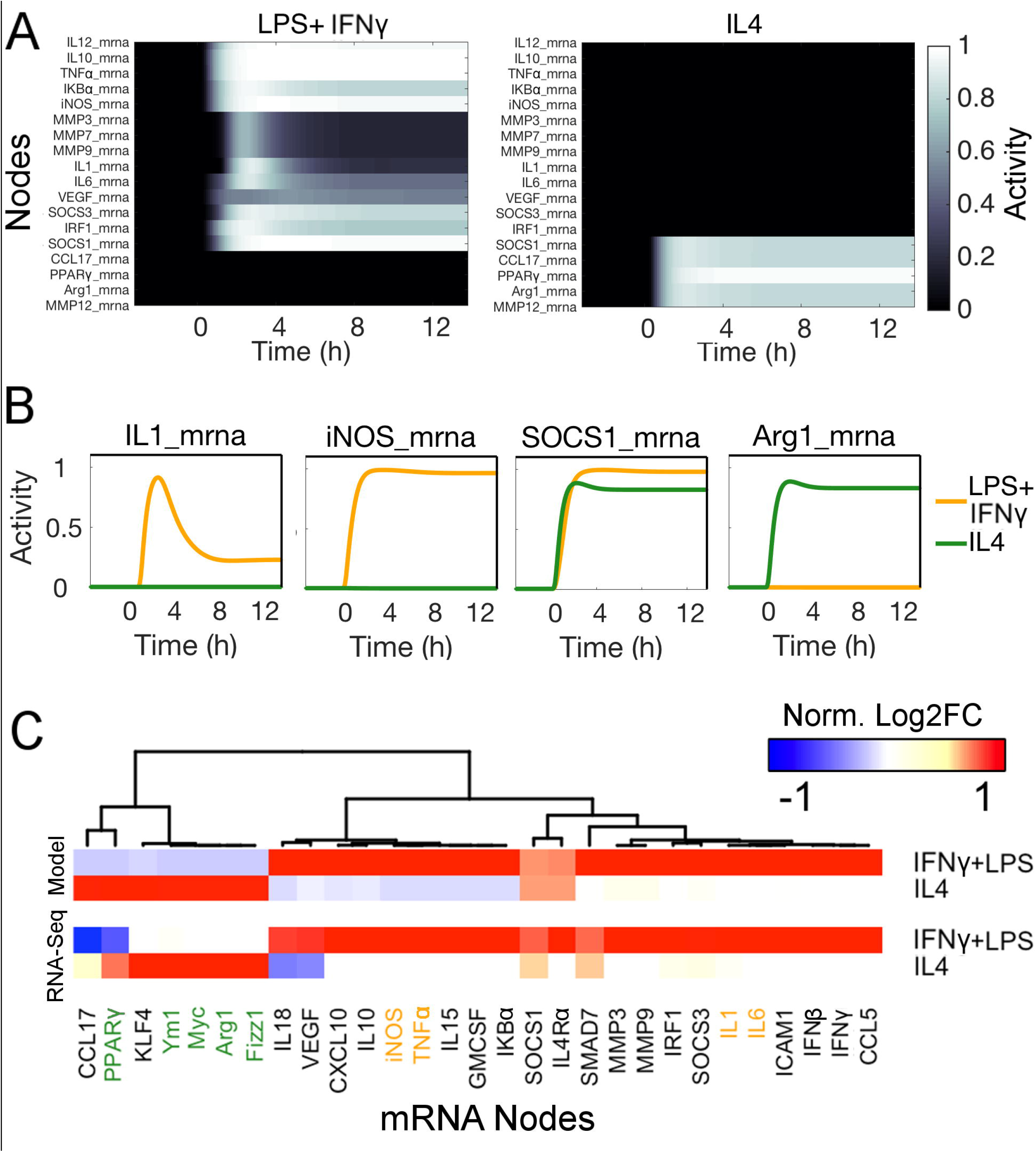
Distinct network dynamics predicted in response to LPS+IFNγ and IL4. A) Dynamics of predicted gene expression in response to stimulation with LPS+IFNγ or IL4. Stimuli were added at 0 h. B) Kinetics of selected mRNAs in response to stimulation with LPS+IFNγ or IL4. C) mRNA expression profiles predicted by the model, validated against RNA-Seq measurements from peritoneal macrophages treated with LPS+IFNγ or IL4 for 4h. For semi-quantitative comparison between model and experiment, the log2 fold change of each mRNA vs. control was normalized by the root mean square between the M1 and M2 conditions. Classic M1 (orange) and M2 (green) phenotype markers are highlighted.

Model predictions of mRNA expression were compared to experimental transcriptome responses of peritoneal macrophages stimulated with LPS+IFNγ or IL4 for 4 h (**Figure 2C**; see **Supplementary Figure S2** for differential expression analysis). Semi-quantitative comparisons between the model and experimental measurements were performed by root-mean squared (RMS) normalization of log2 fold changes in gene expression. For both LPS+IFNγ and IL4 stimulated conditions, we observed high consistency between the predicted model and experimentally measured expression profiles. In the LPS+IFNγ stimulated macrophages, 27 out of 29 genes were semi-quantitatively consistent (absolute difference in RMS-normalized fold change less than 0.4). The two quantitatively inconsistent genes, C-C motif chemokine ligand 17 (CCL17) and peroxisome proliferator-activated receptor-γ (PPARγ), both qualitatively decreased in the RNA-Seq data and model predictions. In IL4-stimulated macrophages, 26 out of 29 genes were semi-quantitatively consistent. Two of the three inconsistent genes, CCL17 and SMAD7, both qualitatively increased in the RNA-Seq data and model prediction. IL4 Receptor-α (IL4Rα) was predicted to be increase yet was not significantly differentially expressed in the RNA-Seq data. Overall the model exhibited 91.4% (53 of 58) semi-quantitative match and another 6.9% (4 of 58) trend match with RNA-Seq data, for a total match of 98.3% (57 of 58).

To identify the key drivers of differential macrophage responses to LPS+IFNγ and IL4 input-dependent differential responses, we simulated network-wide node knockdowns. As shown in **Supplementary Figure S3**, the network response to knockdowns differed considerably between LPS+IFNγ and IL4 conditions. Network influence of a given node was quantified by summing the absolute change in all network nodes when that node was knocked down (columns in **Supplementary Figure S3**). The most influential nodes in LPS+IFNγ-treated macrophages differed considerably from the most highly influential nodes with IL4 treatment (**Figure 3A**). Node sensitivity was quantified by summing the absolute change in that node across all node knockdowns (rows in **Supplementary Figure S3**).

**Figure 3.**
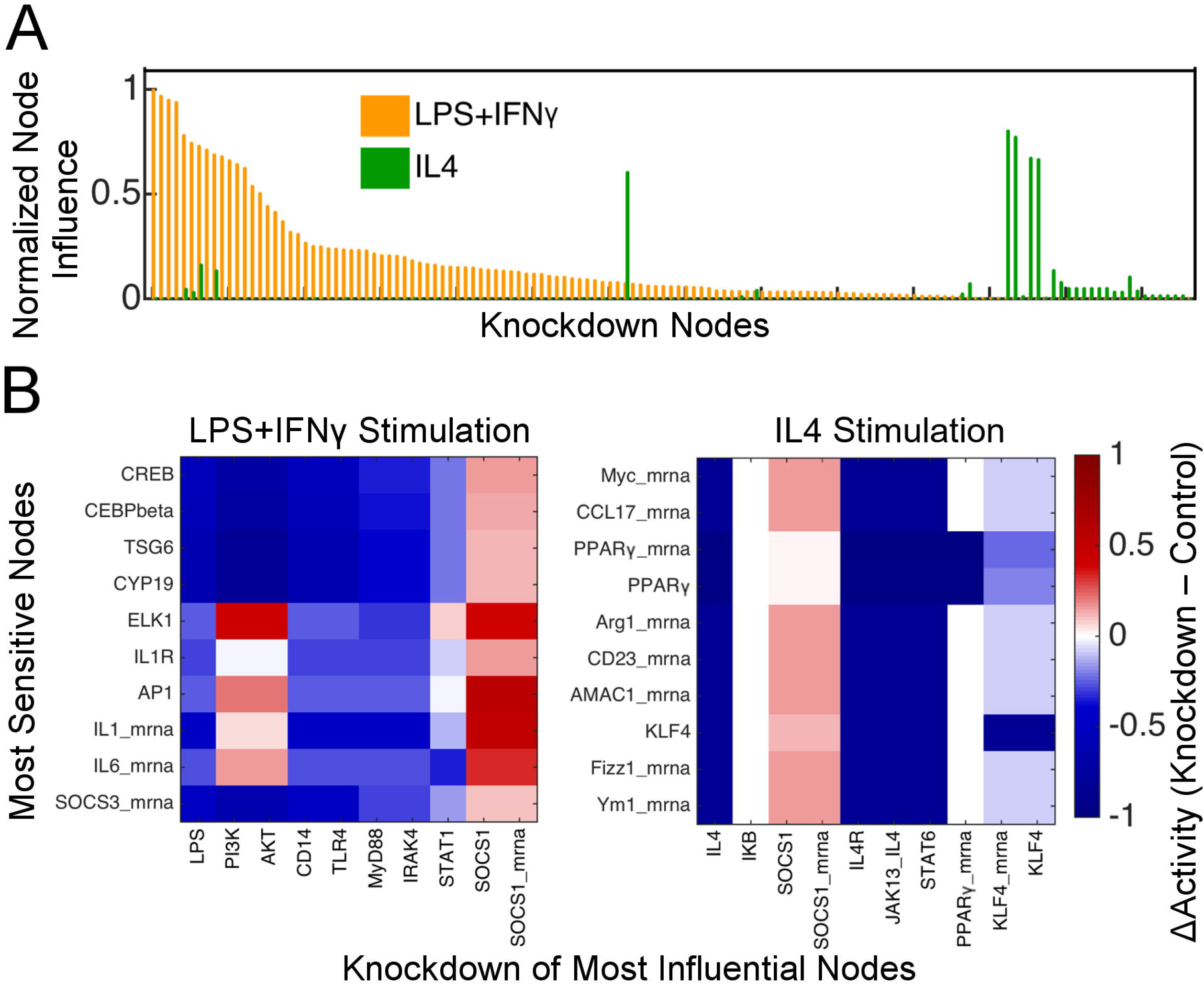
Network-wide knockdown simulations predict distinct mechanistic drivers of macrophage activation with pro- and anti-inflammatory stimuli. A) Overall network influence of node knockdowns under stimulation with either LPS+IFNγ (orange) or IL4 (green). Nodes were ranked by the overall influence of their knockdown on all other network nodes, under conditions of LPS+IFNγ stimulation. B) Predicted effect of knockdown of influential nodes on activity of highly sensitive nodes, under conditions of LPS+IFNγ or IL4 treatment.

Based on network-wide knockdown simulations, the top 10 most influential nodes and top 10 most sensitive nodes were ranked for both the LPS+IFNγ and IL4 stimulated conditions. Under LPS+IFNγ stimulation, the most influential nodes are the LPS-toll like receptor 4 (TLR4)-myeloid differentiation 88 (MyD88)- TNF receptor associated factor 6 (TRAF6) signaling axis, phosphoinositide 3-kinases (PI3K)/AKT, and pro-inflammatory transcriptional factors signal transducer and activator of transcription 1 (STAT1) and nuclear factor κB (NFκB) (**Figure 3B**, **left panel**). The nodes most sensitive to knockdowns under LPS+IFNγ stimulation were induced by mitogen-activated protein kinases (MAPKs) and IL1 autocrine signaling, suggesting a highly interactive and feedback-dependent network. In contrast, with IL4 stimulation the most influential nodes were associated with the IL4-STAT6 signaling axis except IκBα (which was negatively regulated by the IL4-STAT6 pathway) and SOCS1, which negatively fed back to STAT6 activation. The most sensitive nodes under IL4 stimulation were all STAT6-induced, consistent with the dominant signaling through the IL4-STAT6 signaling axis (**Figure 3B**, **right panel**).

### Distinct macrophage phenotypes predicted in response to stimuli combinations

During inflammation, macrophages are subjected to multiple, sometimes conflicting cues. Responses to combinations of stimuli may reveal the crosstalk mechanisms that underlie cellular decision making. To this end, we simulated the 9 single input stimuli, 36 pairwise combinations, and negative control conditions. Network responses to cytokine combinations clustered into 6 phenotypes, which were largely determined by a dominating role of LPS, TNFα/IFNγ/IFNβ, IL1, or IL4 (**Figure 4A**, conditions listed in **Table 1**). Signaling modules distinctly induced by these stimuli are visualized in **Supplementary Figure S4**.

**Figure 4.**
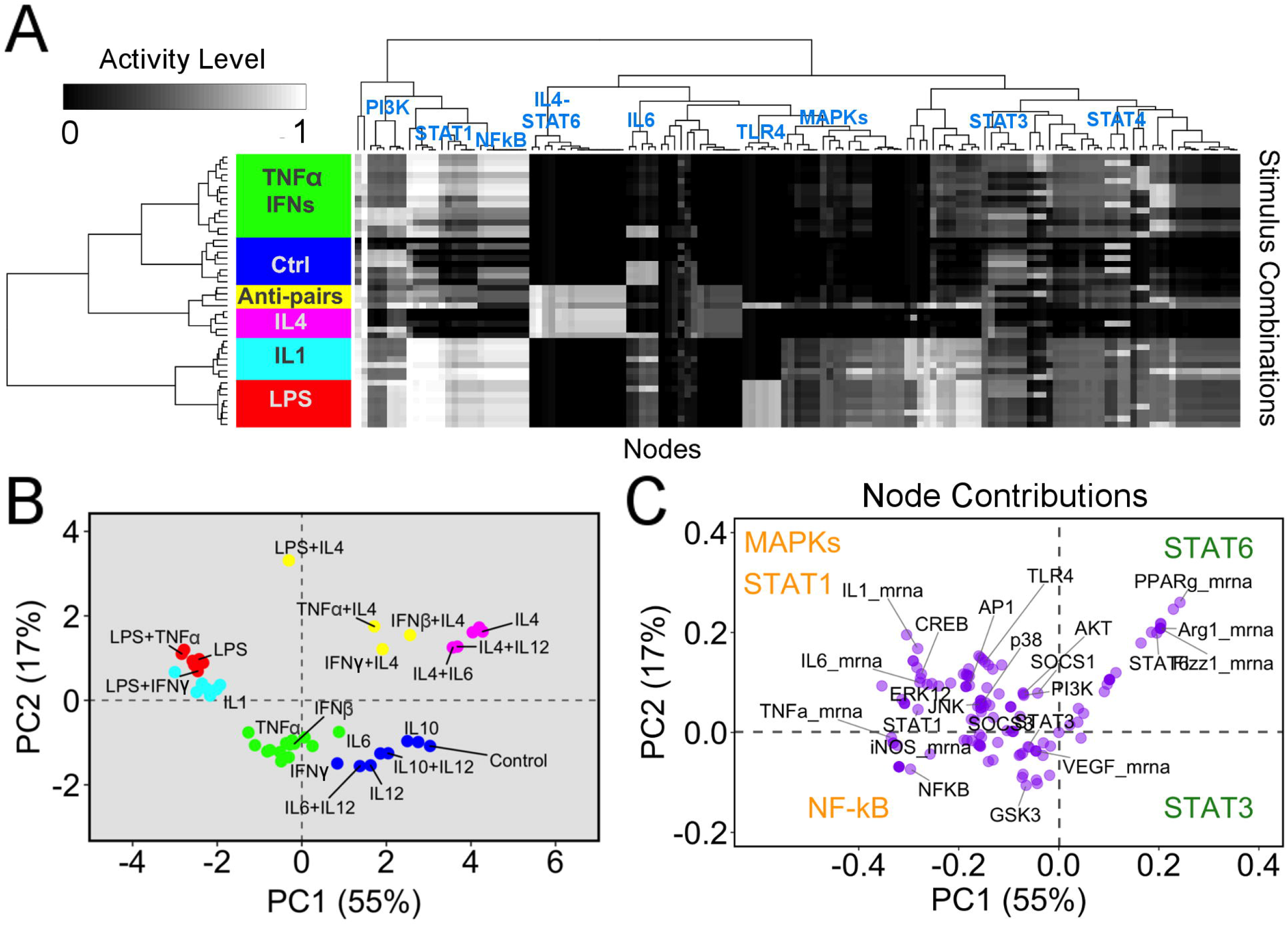
Distinct macrophage activation states induced by combined stimuli. A) Network-wide response to 9 single input stimuli, 36 pairwise combinations, and negative control conditions at 4 h. Hierarchical clustering was performed to identify six phenotype clusters (color coded rows) and signaling modules (column dendrogram). B) Principal component analysis (PCA) scores reveal relationships between the six phenotype clusters induced by combined stimuli. C) PCA loadings indicate the contribution of each node’s activity to the PC1 and PC2 dimensions. M1-associated (orange) and M2-associated (green) labels indicate representative signaling modules within the quadrants.

**Table 1.**
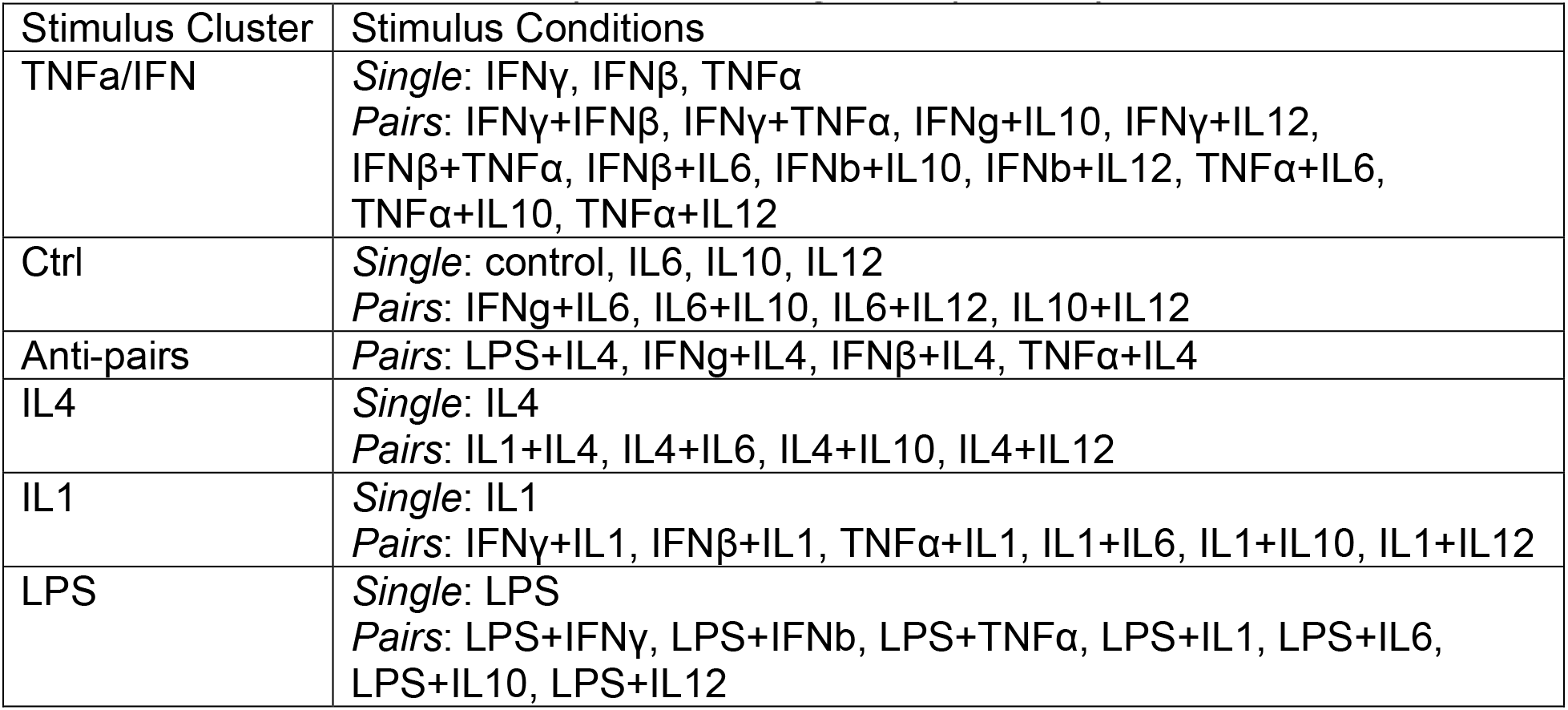
Clusters of network responses to single and paired cytokine stimuli.

| Stimulus Cluster | Stimulus Conditions |
| --- | --- |
| TNFα/IFN | *Single:* IFNγ, IFNβ, TNFα<br>*Pairs:* IFNγ+IFNβ, IFNγ+TNFα, IFNγ+IL10, IFNγ+IL12, IFNβ+TNFα, IFNβ+IL6, IFNβ+IL10, IFNβ+IL12, TNFα+IL6, TNFα+IL10, TNFα+IL12 |
| Ctrl | *Single:* control, IL6, IL10, IL12<br>*Pairs:* IFNγ+IL6, IL6+IL10, IL6+IL12, IL10+IL12 |
| Anti-pairs | *Pairs:* LPS+IL4, IFNγ+IL4, IFNβ+IL4, TNFα+IL4 |
| IL4 | *Single:* IL4<br>*Pairs:* IL1+IL4, IL4+IL6, IL4+IL10, IL4+IL12 |
| IL1 | *Single:* IL1<br>*Pairs:* IFNγ+IL1, IFNβ+IL1, TNFα+IL1, IL1+IL6, IL1+IL10, IL1+IL12 |
| LPS | *Single:* LPS<br>*Pairs:* LPS+IFNγ, LPS+IFNβ, LPS+TNFα, LPS+IL1, LPS+IL6, LPS+IL10, LPS+IL12 |

Principal component analysis separated M1-like phenotypes stimulated by pro-inflammatory cytokines from the M2-like phenotype stimulated by anti-inflammatory cytokines along principal component 1 (PC1) (**Figure 4B**). Principal component 2 (PC2) provided further distinction among macrophage phenotypes beyond the well-established M1-M2 axis. LPS and IFNγ are both considered classic M1-inducing stimuli [1], [31], and they both strongly stimulated the NFκB module. However, LPS was distinguished along PC2 by stronger activation of MAPKs and STAT1 modules and IL1 mRNA expression, while IFNγ stimulated glycogen synthase kinase 3 (GSK3) (**Figure 4C**). IL4- and IL10-dominated combinations were both located in the positive PC1 direction, associated with a M2-like phenotype. However, PC2 distinguished their distinct regulation of STAT6 and STAT3 modules (**Figure 4C**), which is consistent with previously reported distinctions between M2-like phenotypes induced by IL4 and IL10.[38], [39] IL10-treated macrophages are generally considered as a deactivated M2 phenotype, consistent with the IL10-induced phenotypes clustered together with the control condition.

Co-stimulation of LPS, TNFα, IFNγ, or IFNβ with IL4 produced a mixed phenotype distinct from that observed with any individual stimulus (**Figure 4B**). As expected, combinations of these pro- and anti-inflammatory stimuli were mutually inhibiting along the M1-M2 axis. Surprisingly, these combinations were mutually activating along the PC2 dimension. PCA did not resolve unique markers of the mixed phenotype, indicating that closer examination of particular conflicting stimuli was needed to identify the drivers of mutual inhibition and activation. Compared to analysis of single treatments alone (**Supplementary Figure S5**), combination treatments decreased the variance explained by PC1 from 61% to 55% and increased the variance explained by PC2 from 14% to 17%. Together these results indicate an important dimension to macrophage activation beyond the classic M1-M2 polarization paradigm.

### Antagonistic stimulus combinations elicited both antagonistic and mutualistic responses in different signaling modules

To identify network mechanisms that may contribute to cross-talk between conflicting cues, we focused on IFNγ with IL4, as this pair often co-exists in vivo and has been studied experimentally.[40] Signaling module activation was quantified by the sum of the node activities within each module, as identified in the hierarchical clustering analysis. Addition of a conflicting stimulus decreased activity of IFNγ-induced MAPKs, NFκB, and STAT1 modules and IL4-induced STAT6 modules, demonstrating mutual inhibition of these modules (**Figure 5A**). In contrast, STAT3 and PI3K modules were further activated by co-stimulation with the pro- and anti-inflammatory inputs, consistent with our observation of a unique mixed phenotype.

**Figure 5.**
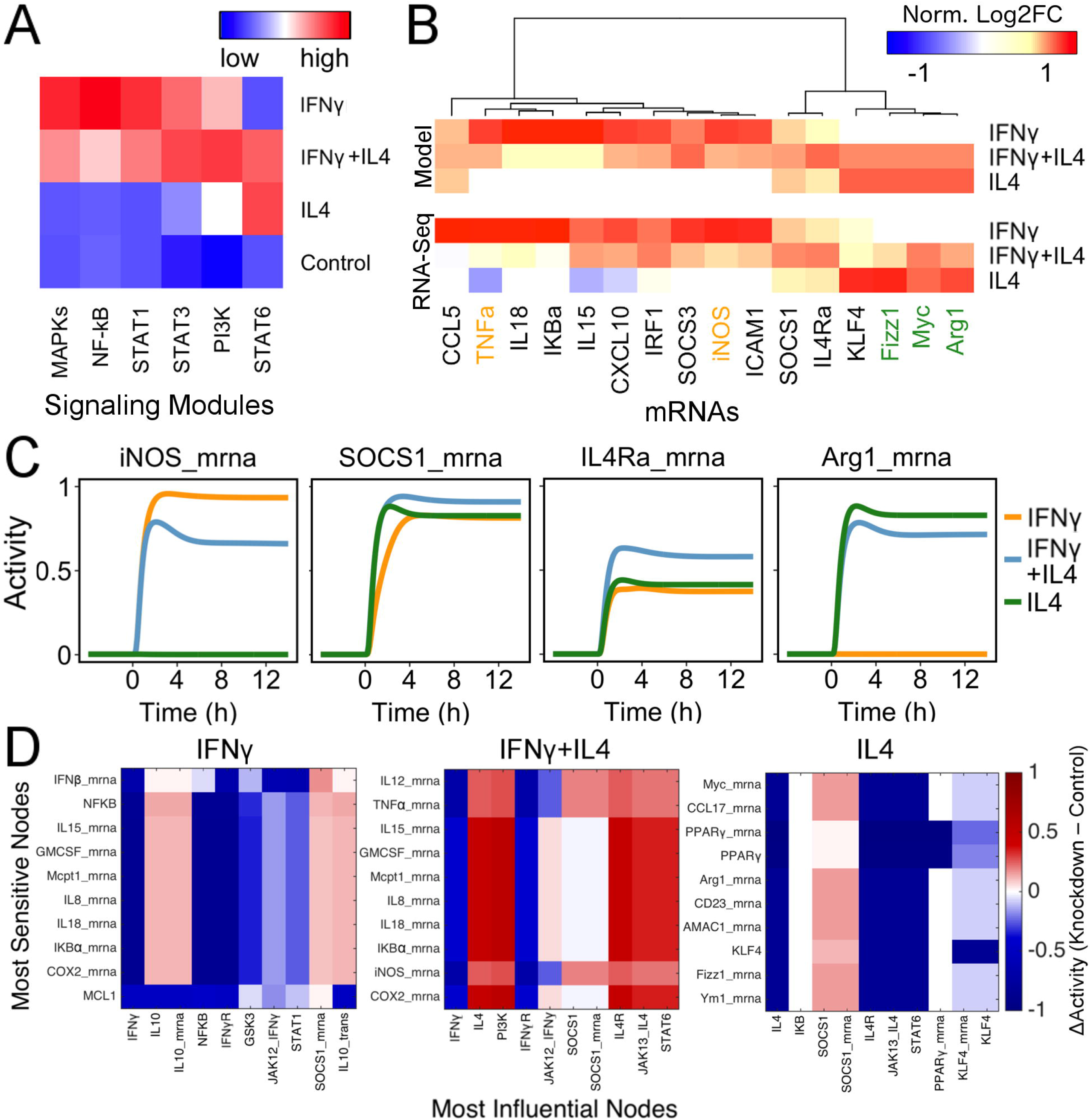
Macrophage network model predicts both mutual inhibition and mutual activation in response to conflicting cues. A) Model-predicted signaling module activities in response to IFNγ and IL4 treatments at 4h, column normalized. B) Experimental validation of mRNA expression predicted in response to IFNγ, IL4, or IFNγ+IL4. For both experimental data [40] and model predictions, mRNA were independently normalized by RMS-normalized log2 fold change at 4 h. C) Predicted expression dynamics of selected mRNAs in response to IFNγ, IL4, or IFNγ+IL4. D) Context-dependent network response to node knockdowns under treatments of IFNγ, IL4, or IFNγ+IL4.

We further examined potential cross-talk between IFNγ and IL4 on gene expression, which was validated against independent published RNA-Seq data of murine bone marrow-derived macrophages treated with IFNγ and IL4 combinations for 4 h (**Figure 5B**).[40] The difference in RMS-normalized change in gene expression between single cytokine (IFNγ or IL4) and combined IFNγ+IL4 conditions was computed for both model and experiments. Genes were grouped as IFNγ-, IL4-, or mutually-induced based on the RNA-Seq responses. The model correctly predicted nine IFNγ-induced genes suppressed by co-stimulation with IL4 (TNFα, IL18, IKBα, IL15, CXCL10, IRF1, SOCS3, iNOS, ICAM1). One exception was CCL5 mRNA, which was not predicted to be differentially regulated by either IFNγ or IL4. The model also correctly predicted IFNγ-mediated inhibition of four IL4-induced genes (KLF4, Fizz1, Myc, Arg1). In addition to these mutually inhibitive effects, the model correctly predicted mutual induction of SOCS1 and IL4Rα gene expression by IFNγ+IL4 co-stimulation (predicted kinetics shown in **Figure 5C**).

Responses to IFNγ+IL4 co-stimulation were visualized to identify network mechanisms contributing to mutual inhibition or activation (**Supplementary Figure S6**). SOCS1 mRNA was induced by IFNγ-stimulated interferon regulatory factor 1 (IRF1) and IL4-stimulated STAT6. Mutual induction of IL4Rα mRNA was mediated by IFNγ-stimulated STAT3 and IL4-stimulated STAT6. Mutual activation of PI3K/AKT was mediated by IFNγ-stimulated IL10 and TNFα as well as IL4-stimulated growth factor receptor-bound protein 2 (GRB2). Under combined IFNγ+IL4, network-wide knockdowns demonstrate that mutually activated PI3K and SOCS1 became highly influential in suppressing pro-inflammatory and anti-inflammatory genes, respectively (**Figure 5D** and **Supplementary Figure S7**).

## Discussion

Here, we developed a computational model that provides a quantitative framework with which to understand how macrophages integrate and respond to multiple, often conflicting cues. The model was validated against transcriptome measurements from pro- and anti-inflammatory cues (LPS+IFNγ and IL4, respectively), as well as mixed IFNγ + IL4 stimulation. In response to combined treatments, macrophages were predicted to respond not only along the classic M1-M2 polarization axis but also along a second, orthogonal dimension differentiating inactive (M0) macrophages from macrophages that are activated by mixed cues. The model predicted key network mechanisms that mediate mutual inhibition among M1 and M2-associated cues, which include predicted mutual activation of the PI3K/STAT3 signaling module and enhanced gene expression of SOCS1 mRNA and IL4Rα. Overall, this study illustrates how systems analysis of responses to combined stimuli can reveal network principles that underlie cellular decision making.

The classic M1-M2 paradigm distinguishes between pro- and anti-inflammatory macrophages through differential expression of phenotype markers (*e.g.* IL1, IL6, iNOS, TNFα for M1; (Arg1, found in inflammatory zone 1 (Fizz1), PPARγ for M2).[1], [3], [31] *In vitro* studies frequently use LPS, IFNγ, or LPS+IFNγ treatment to induce the M1-like phenotype and IL4 or IL10 to induce the M2-like phenotype in mouse, although each stimulus yields a somewhat different activation state. Furthermore, these simplified stimulation conditions do not replicate the dynamic multi-factorial stimuli macrophages experience in vivo.[1], [2], [20], [21] The signaling network mediating macrophage activation is highly complex, making comprehensive perturbations of cytokine and intracellular perturbations experimentally intractable.[15], [40], [41]

Model predictions of response to classic M1/M2 polarization stimuli LPS+IFNγ or IL4 were largely consistent with RNA sequencing data from peritoneal macrophages and predicted distinctly influential signaling nodes under these conditions. In response to 36 stimulus pairs, the macrophage network model responded not only along the classic M1-M2 polarization axis but also along a second axis that further differentiated among macrophage phenotypes (**Figure 6**). Along this new dimension, antagonistic combinations of IL4 and IFNγ or other pro-inflammatory stimuli (LPS induced a mixed phenotype distinct from either inactive or M1/M2 polarized macrophages. Many classic M1 and M2 markers were mutually inhibited, yet the PI3K signaling module and SOCS1 and IL4Rα mRNAs (in the STAT3 module) were mutually activated. Knockdown simulations predicted that SOCS1 and PI3K were not only responsive but also helped to mediate the mutual inhibition characteristic of the mixed phenotype.

**Figure 6.**
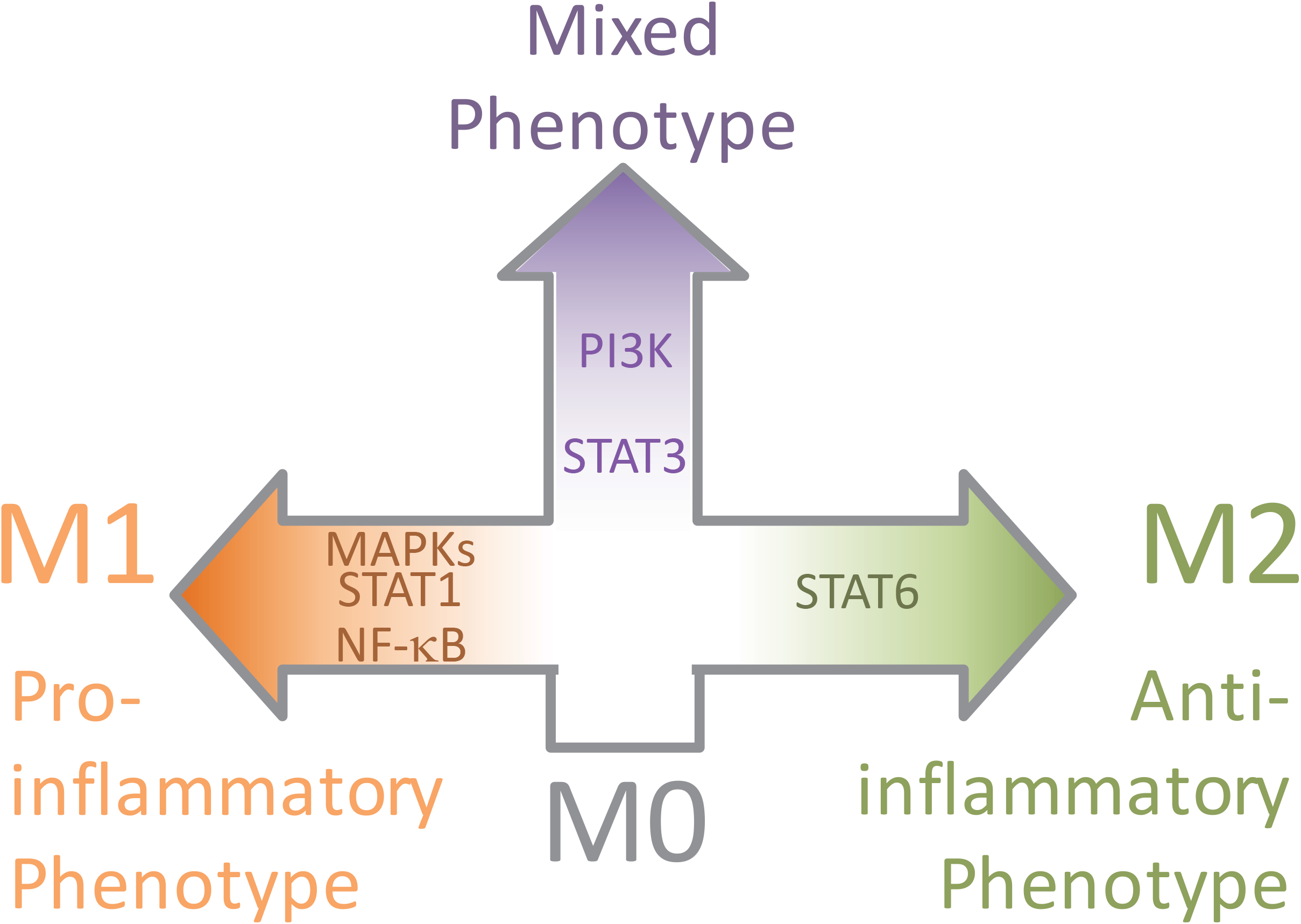
Illustrative roadmap of macrophage activation phenotypes and signaling modules induced by combinations of stimuli. Combinations of pro- and anti-inflammatory stimuli induced a distinct mixed phenotype associated with mutual activation of PI3K and STAT3 modules yet mutual inhibition of M1- and M2- associated markers.

Macrophage phenotypes are typically assessed based on markers of mRNA or protein abundance.[1], [3], [31] Here, modeling of dynamic post-translational regulation of signaling increased the ability to resolve distinct macrophage phenotypes, particularly in response to antagonistic cytokine combinations and at early timepoints. PI3K, AKT and GSK3 activities were among the strongest contributors to the mixed phenotype activation axis orthogonal to the M1-M2 polarization axis. Simulated knockdown of PI3K and AKTt were also highly influential on macrophage network state with combined IFNγ + IL4 stimulation. Indeed, PI3K/AKT signaling has been described as a converging point for macrophage activation in response to multiple inflammatory stimuli, with distinct roles depending on the stimulus.[42] Further, signaling dynamics aided understanding of the mechanisms by which mRNA markers of macrophage phenotype were mutually inhibited or stimulated.

Macrophage activation has previously been modeled using alternative modeling formalisms and differing scope. A previous Boolean model of macrophage polarization was developed and specifically refined based on data from bone marrow-derived macrophages treated with LPS or IL4 with IL13 [32]. For the 10 genes in common between their model and ours, a logic-based differential equation version of the Boolean model also predicts gene expression in response to LPS or IL4 that is mostly consistent with the RNA-Seq data from peritoneal macrophages used in our analysis (**Supplementary Figure S8**). The model developed here incorporates additional cytokine inputs (IFNγ, IL1, IL6, IL12, and TNFα), cross-talk mechanisms, and genes that were important for analysis of combined stimuli. The logic-based differential equation framework allowed prediction of continuous dynamics and levels of all nodes in the network, allowing semi-quantitative comparisons of perturbation responses, experimental validation, and future analysis of dynamically varying inputs. While we focused analysis towards early changes in signaling and transcription, models focusing on downstream kinetics of gene regulation have shown an important role for regulation of mRNA stability.[43] Future model revisions incorporating mRNA stability,[43] microRNAs,[32] and chromatin modifications[40] may provide further insight into the feedbacks guiding macrophage activation dynamics.

In conclusion, the macrophage network model developed here provides a framework for network-based understanding of how macrophages respond to complex stimuli. Integrated network analyses and experimental studies in the context of mixed stimuli are needed to better characterize and understand the spectrum of macrophage phenotypes in physiologic and pathologic settings.

## Materials and Methods

### Model development

An initial macrophage signaling network was constructed based on literature search in PubMed, identifying review articles and original articles using the search terms “macrophage polarization”, “macrophage activation”, “computational modeling”, and “peritoneal macrophages”.[1], [30]–[32] The signaling network was then extended to include additional established macrophage activation markers that were differentially expressed in peritoneal macrophages from wild type (WT) C57/BL6J mice treated with either 1 μg/ml LPS and 20 ng/ml IFNγ or 20 ng/ml IL4 for 4h (see *RNA-seq analysis*, below).

Differences between initial model predictions and experimental measurements indicated an important role of crosstalk between pro- and anti-inflammatory stimuli. This motivated further model extension through focused literature search on 1) autocrine loops identified with keywords such as “macrophage signaling pathway” IFNβ, IL10, or IL12;[44]–[49] 2) inclusion of feedback loops reported for SOCS and GSK3;[50], [51, p. 1], [52, p. 3], [53], [54] and 3) the addition of key nodes such as PI3K and cAMP response element-binding protein (CREB).[15], [30], [48], [55] Autocrine loop candidates were first identified by reviewing the significantly induced cytokines in the LPS+IFNγ and IL4 stimulated macrophages, indicating roles for the IL12-STAT4 and the IL10-STAT3 signaling axes. The core feedback nodes including SOCS1, SOCS3, GSK3 were examined next and added. Additional signaling modules reported as key cross-talking nodes of multiple pathways such as PI3K and CREB were also added into the network. The finalized macrophage signaling network model includes 9 cytokines critical in macrophage polarization, LPS, IFNγ, IFNβ, IL1, IL4, IL6, IL10, IL12, and TNFα. The model consists of 139 nodes (mRNA, proteins, and small molecules) and 200 reactions.

The signaling network structure was automatically translated into a logic-based differential equation model as previously described[27], [36], [37], [56] using open source Netflux software (https://github.com/saucermanlab/Netflux). The activity of each node was modeled using ordinary differential equations with steady state properties determined by normalized Hill activation or inhibition functions with default parameters and continuous AND/OR logic gating[56]. Default reaction parameters include reaction weight (1), Hill coefficient (1.4), and EC50 (0.5). Default node parameters include yinit (0) and ymax (1).[56] The node parameter τ (time constant) was scaled according to the type of node: 6 min for signaling post-translational modifications, 30 min for mRNA expression, and 1 h for protein expression based on previous macrophage-specific studies.[43], [57]–[60] Reactions weights involving protein translation, or with multiple inputs were set to 0.5 to avoid basal saturation. The baseline level of input was defined as 5% activity for all inputs (weight = 0.05). Where specified, simulations of particular cytokine stimuli (LPS+IFNγ or IL4) were performed by increasing the weights of corresponding input reactions from 5% to 70% (weight = 0.7). The model was simulated in MATLAB v2015b using the adaptive time step solver ODE15S.

### RNA-seq analysis and semi-quantitative model validation

All animal procedures were approved by the Institutional Animal Care and Use Committee at the University of Mississippi Medical Center and were conducted in accordance with the Guide for the Care and Use of Laboratory Animals published by the United States National Institutes of Health (Eighth edition; revised 2011). Peritoneal macrophages were isolated from adult (3-6 month old) C57BL/6J mice (n=4) as previously described.[61], [62] Cells were plated at 1.5×10^6^ cells/well, incubated overnight at 37°C, and then washed with fresh media. Macrophages were assigned to one of three treatment groups: 1) stimulated with 1 μg/mL LPS (Sigma, L2880) and 20 ng/mL IFNγ (R&D, 485-MI) for 4 h; 2) stimulated with 20 ng/mL IL4 (R&D, 404-ML) for 4 h; or 3) untreated for 4 h, serving as the negative control.

Transcriptome measurements and analyses were performed as previously described [63], [64]. RNA was extracted using the Pure Link RNA Mini Kit (Ambion, Foster City, CA) in accordance with manufacturer instructions. cDNA libraries were assembled using the TruSeq Total Stranded RNA with RiboZero Kit (Ambion), set-A, quantified using the Qubit System (Invitrogen, Carlsbad, CA). cDNA library size and quality were determined with the Experion DNA 1K Chip (Bio-Rad, Hercules, CA). cDNA libraries were sequenced using the NextSeq 500 High Output Kit (300 cycles, paired end 100 bp) on the Illumina NextSeq 500 platform (Illumina, San Diego, CA). Sequenced reads (length = 30–50; Cloud Computing Platform), and Fastq file sequences were aligned to the reference genome USCS-GRCm38/mm10 using the STAR aligner with the RNA-Seq Alignment Application [65]. RNAseq count matrices were analyzed for differential mRNA expression compared to the untreated group (adjusted p-value < 0.05) using the R ‘DESeq2’ package [66]. IL4-treated and LPS+ IFNγ-treated groups were each separately compared to the untreated group. Gene set enrichment analysis was performed with Reactome2016 pathways in EnrichR, which uses Fisher’s exact test to compute a combined score as c = ln(p-value)*(z-score) [67]. For heatmap visualization, normalized counts output from DESeq2 were normalized by log10(counts per million). All statistical analysis was performed using R version 3.5.1 and RStudio 1.0.143.

For comparison to model predictions, experimentally measured log2 fold changes of mRNA compared to negative control were normalized by the root mean square (RMS) between treatment groups. Likewise, model-predicted log2 fold changes in mRNA compared to negative control (baseline inputs 5%) were normalized by the RMS between treatment groups. Genes were classified as semi-quantitatively consistent if the absolute difference between model and experimentally measured RMS-normalized log2 fold change was smaller than 0.4 (20% of the ±1 range).

Published RNA-Seq data of murine bone marrow-derived macrophage (BMDMs) treated with IFNγ, IL4, IFNγ+IL4, or negative control for 4 h [40] were obtained from Gene Expression Omnibus with the *GEOquery* package in R (GSE84520). *DESeq2* [66] was applied to identify differentially expressed genes (adjusted p-value < 0.05). These data were used as a second validation of IL4 predictions, as well as to validate predictions of IFNγ and combined IFNγ+IL4. The difference in RMS-normalized change in gene expression between single cytokine (IFNγ or IL4) vs. combined IFNγ+IL4 conditions was computed for both model and experiments.

### Sensitivity analysis

Comprehensive single-knockdowns were simulated to identify the functional influence of each node in a given experimental condition.[37] Complete knockdown was simulated by setting ymax = 0 for that node. Change in activity was calculated as the difference in an individual node activity with and without knockdown in response to the specified stimulus at 4 h. The sensitivity of a node in a given condition was quantified by summing the absolute activity changes for that node across all node knockdowns (e.g. the corresponding row of **Supplementary Figure 3**). The influence of a node in a given condition was quantified by summing the absolute activity changes of all nodes in response to that knockdown of that node (e.g. the corresponding column of **Supplementary Figure 3**).

### Combined stimuli screening

Network responses to the 9 single inputs, 36 pairwise combinations, and control conditions were hierarchically-clustered to identify macrophage phenotypes and signaling modules. Phenotypes were identified by clustering across conditions (rows) using the Ward method, focusing on the variance between different treatment responses. Signaling modules were identified by clustering across nodes (columns) using the complete linkage method, which focuses on the associations among the different signaling nodes. Module activities were calculated as the sum of node activities within each module. Principal component analysis (PCA) and variable contribution analysis was performed using the FactoMineR package in R.

## Authorship

Conceptualization: JJS, ML; Investigation: XL, JZ, ACZ; Data curation-Formal analysis: ARN; Writing-original draft: XL, JZ; Writing-editing and revision: XL, ARN, MLL, JJS.

## Acknowledgments

This study was supported by grants from the National Institutes of Health (HL137755, HL127944, HL075360, HL129823, and HL137319), the Biomedical Laboratory Research and Development Service of the Veterans Affairs Office of Research and Development under Award Number 5I01BX000505, and the National Science Foundation (1252854).

## Conflicts-of-Interest Disclosure

The authors declare no conflict of interest.

