## Supplementary Materials for "Network analysis reveals a distinct axis of macrophage activation in response to conflicting inflammatory cues"

### Supplementary Materials for
### Network analysis reveals a distinct axis of macrophage activation in response to conflicting inflammatory cues

Xiaji Liu[1], Jingyuan Zhang[1], Angela C. Zeigler[1], Anders R. Nelson[1], Merry L. Lindsey[2], and Jeffrey J. Saucerman[1*]

**Supplementary Figure S1. Distinct macrophage activation network states induced by LPS+IFNγ and IL4.** Each node is annotated with a bar graph that shows that node's normalized activity in response to LPS+IFNγ at 4 h (red bar) and IL4 at 4 h (blue bar).

**Supplementary Figure S2. Peritoneal macrophage gene expression in response to LPS+IFNγ or IL4.** A) Log10CPM-normalized read counts of the 500 genes with highest variance, sorted by hierarchical clustering (n=4 per treatment condition). B) Principal component analysis of macrophage gene expression, including all transcripts. C) Venn diagram of differentially expressed transcripts in LPS+IFNγ or IL4 treatment groups compared to control. D) Gene set enrichment analysis for transcripts with increased or decreased expression in response to LPS+IFNγ or IL4. E) Volcano plots of differential gene expression analysis of LPS+IFNγ-treated and IL4-treated macrophages, annotating selected genes from the macrophage network model.

**Supplementary Figure S3. Network-wide knockdown simulations under conditions of A) LPS+IFNγ and B) IL4.** In each column, the given node was knocked down by setting its ymax parameter to zero. Row shows the change in node activity in response to given knockdowns (columns). Change in activity was calculated as the difference in a node's activity with and without knockdown in response to the specified stimulus (70%) at 4 h. Network influence of a given node was quantified by summing the absolute change in all network nodes when that node was knocked down (columns). Node sensitivity was quantified by summing the absolute change in that node across all node knockdowns (rows). Influence and sensitivity calculations were used to rank nodes for Figure 3.

**Supplementary Figure S4. Signaling modules distinctly regulated by combinatorial stimuli.** Modules were determined by hierarchical clustering of columns in Figure 3A using the complete linkage method. After identifying 24 initial clusters, some clusters were further condensed manually based on their very similar responses and topological similarity (MyD88-dependent IL1R and TLR4; STAT6-dependent mRNA with proteins; MAPK pathways and MMP mRNAs).

**Supplementary Figure S5. Network responses to all single stimuli.** *A)* Network-wide response to 9 single input stimuli and negative control conditions at 4 h. As in Figure 3, hierarchical clustering of stimuli (rows) was performed using the Ward method, while nodes (columns) were clustered using complete linkage. *B)* Principal component analysis (PCA) scores reveal relationships between phenotype clusters induced by single stimuli, color-coded based on clustering from Figure 3A. *C)* PCA loadings indicate the contribution of each node's activity to the PC1 and PC2 dimensions. Orange (M1-associated) and green (M2-associated) labels indicate representative signaling modules within the quadrants.

**Supplementary Figure S6. Distinct macrophage activation network state induced by IFNγ, combined IFNγ+IL4, and IL4.** Each node in the network is color-coded by its activation in response to IFNγ (left third), IFNγ+IL4 (middle third), and IL4 (right third) at 4 h.

**Supplementary Figure S7. Network-wide knockdown simulations under conditions of A) IFNγ and B) IFNγ+IL4.** In each column, the given node was knocked down by setting its ymax parameter to zero. Row shows the change in node activity in response to given knockdowns (columns). Change in activity was calculated as the difference in a node's activity with and without knockdown in response to the specified stimulus (70%) at 4 h. Influence (absolute sum of rows) and sensitivity (absolute sum of columns) calculations were used to rank nodes for Figure 5D.

**Supplementary Figure S8. Comparison of a previously reported macrophage signaling network structure to experimental data.** Rex et al. recently developed a Boolean model of bone-marrow derived macrophage (BMDM) activation in response to LPS and IL4/IL13 stimuli [32]. To evaluate the predictive performance of that network in a manner comparable to our validations performed in Figure 2C, we converted that BMDM model to our logic-based differential equation framework. mRNA expression was examined for 10 genes that overlap between the Rex et al. model, our peritoneal macrophage model, and the peritoneal macrophage RNA-seq data used for validation in Figure 2C (LPS+IFNγ or IL4 for 4h [34]). Out of the 10 genes, there are 9 semi-quantitative matches between the BMDM model predictions and experimental measurements under LPS+IFNγ stimulation, 8 matches under IL4 stimulation, and 1 additional trend match in the inconsistent genes. The overall consistency of 85% (17 in 20) is similar to our model of 91.4% (53 in 58), which has a greater overall scope and incorporates additional genes.

**Supplementary Table S1.** Macrophage signaling network model. This spreadsheet contains the model network structure, parameters, and supporting literature references. This spreadsheet can be opened by Netflux for network simulations.

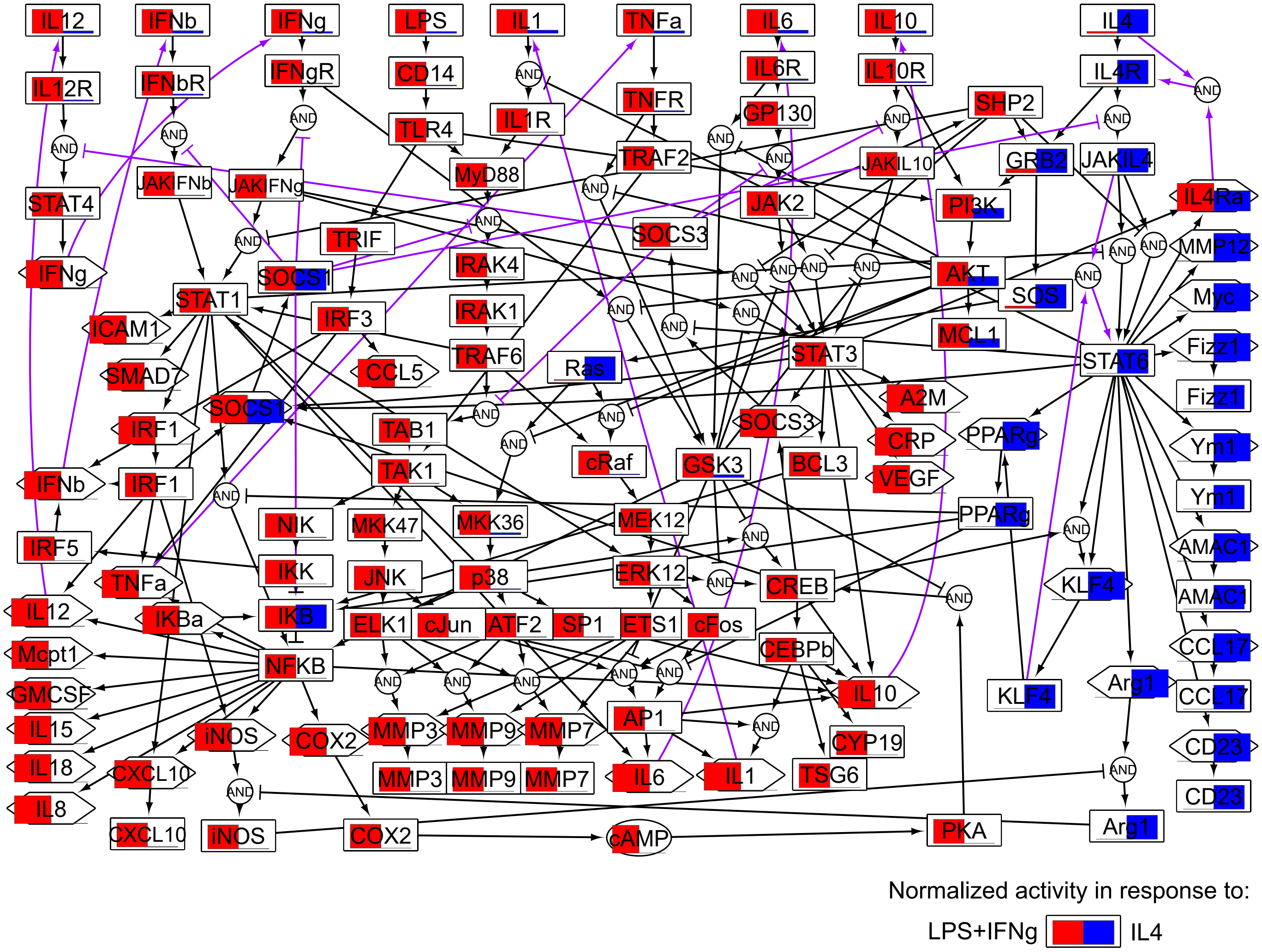
IL12
IFNb
IFNg
LPS
IL1
TNFa
IL6
IL10
IL4
IL12R
IFNbR
IFNgR
CD14
IL6R
IL10R
IL4R
TNFR
GP130
TLR4
IL1R
SHP2
TRAF2
MyD88
JAKIFNb
JAKIFNg
JAKIL10
GRB2
JAKIL4
STAT4
JAK2
PI3K
IL4Ra
TRIF
SOCS3
MMP12
IFNg
IRAK4
SOCS1
AKT
SOS
Myc
STAT1
IRAK1
ICAM1
IRF3
MCL1
Fizz1
STAT3
STAT6
TRAF6
Ras
SMAD
CCL5
A2M
Fizz1
SOCS1
SOCS3
TAB1
IRF1
PPARg
CRP
Ym1
cRaf
GSK3
BCL3
TAK1
VEGF
IFNb
IRF1
Ym1
PPARg
NIK
MKK47
MKK36
MEK12
IRF5
AMAC1
IKK
JNK
p38
ERK12
CREB
TNFa
KLF4
AMAC1
IL12
IKBa
IKB
ELK1
cJun
ATF2
SP1
ETS1
cFos
Mcpt1
CEBPb
CCL17
NFKB
GMCSF
IL10
KLF4
Arg1
CCL17
AP1
IL15
iNOS
COX2
MMP3
MMP9
MMP7
CYP19
CD23
IL18
CXCL10
MMP3
MMP9
MMP7
IL6
IL1
TSG6
IL8
CD23
CXCL10
iNOS
COX2
cAMP
PKA
Arg1
AND
Normalized activity in response to:
LPS+IFNg
IL4

Figure S1

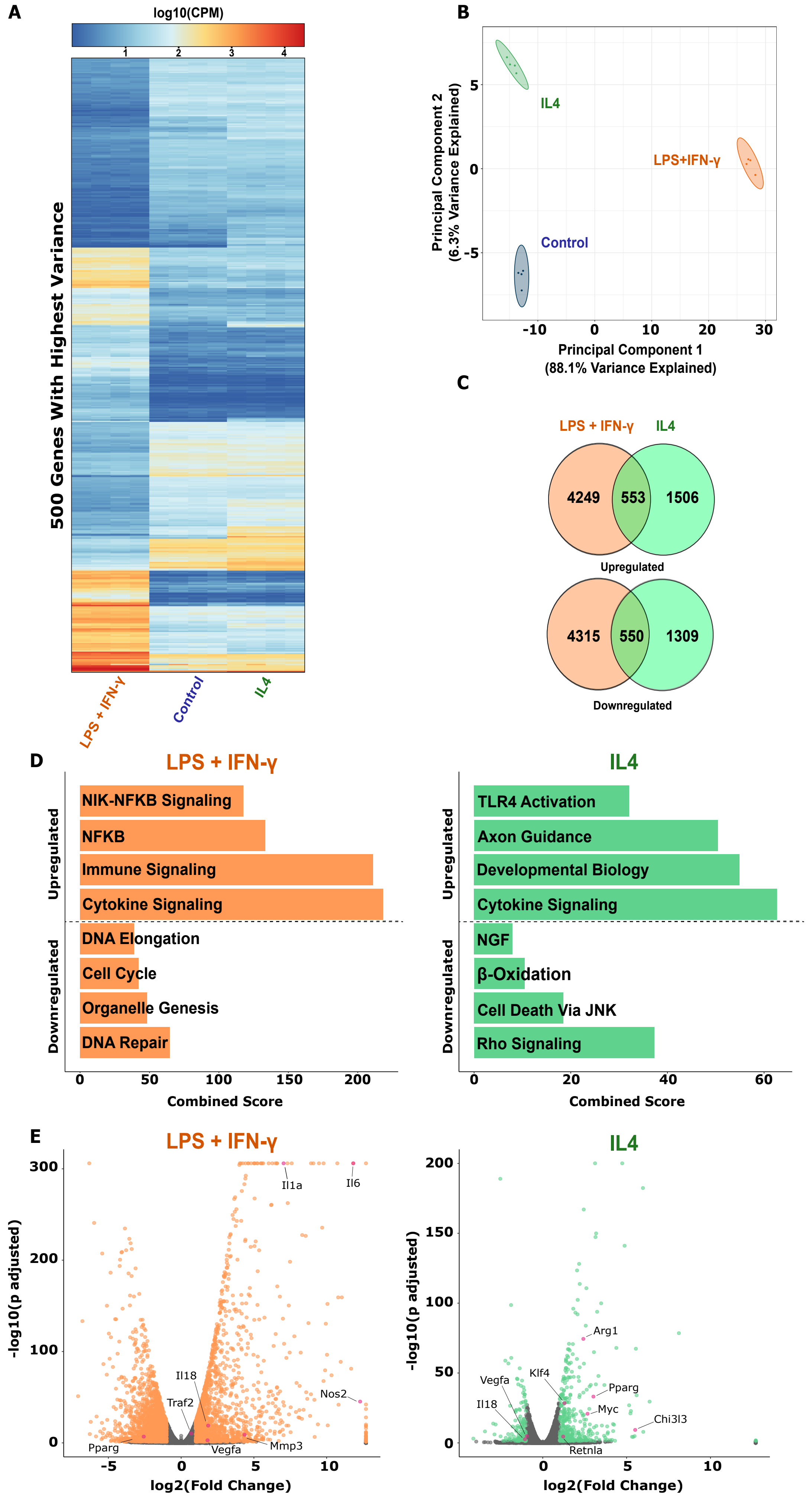
A
log10(CPM)
1
2
3
4
500 Genes With Highest Variance
LPS + IFN-γ
Control
IL4
B
Principal Component 2
(6.3% Variance Explained)
5
0
-5
IL4
LPS+IFN-γ
Control
-10
0
10
20
30
Principal Component 1
(88.1% Variance Explained)
C
LPS + IFN-γ
IL4
4249
553
1506
Upregulated
4315
550
1309
Downregulated
D
LPS + IFN-γ
Upregulated
NIK-NFKB Signaling
NFKB
Immune Signaling
Cytokine Signaling
Downregulated
DNA Elongation
Cell Cycle
Organelle Genesis
DNA Repair
0
50
100
150
200
Combined Score
IL4
Upregulated
TLR4 Activation
Axon Guidance
Developmental Biology
Cytokine Signaling
Downregulated
NGF
β-Oxidation
Cell Death Via JNK
Rho Signaling
0
20
40
60
Combined Score
E
LPS + IFN-γ
-log10(p adjusted)
300
200
100
0
Il1a
Il6
Il18
Traf2
Nos2
Pparg
Vegfa
Mmp3
-5
0
5
10
log2(Fold Change)
IL4
-log10(p adjusted)
200
150
100
50
0
Arg1
Klf4
Vegfa
Pparg
Il18
Myc
Chi3l3
Retnla
0
5
10
log2(Fold Change)

Figure S2

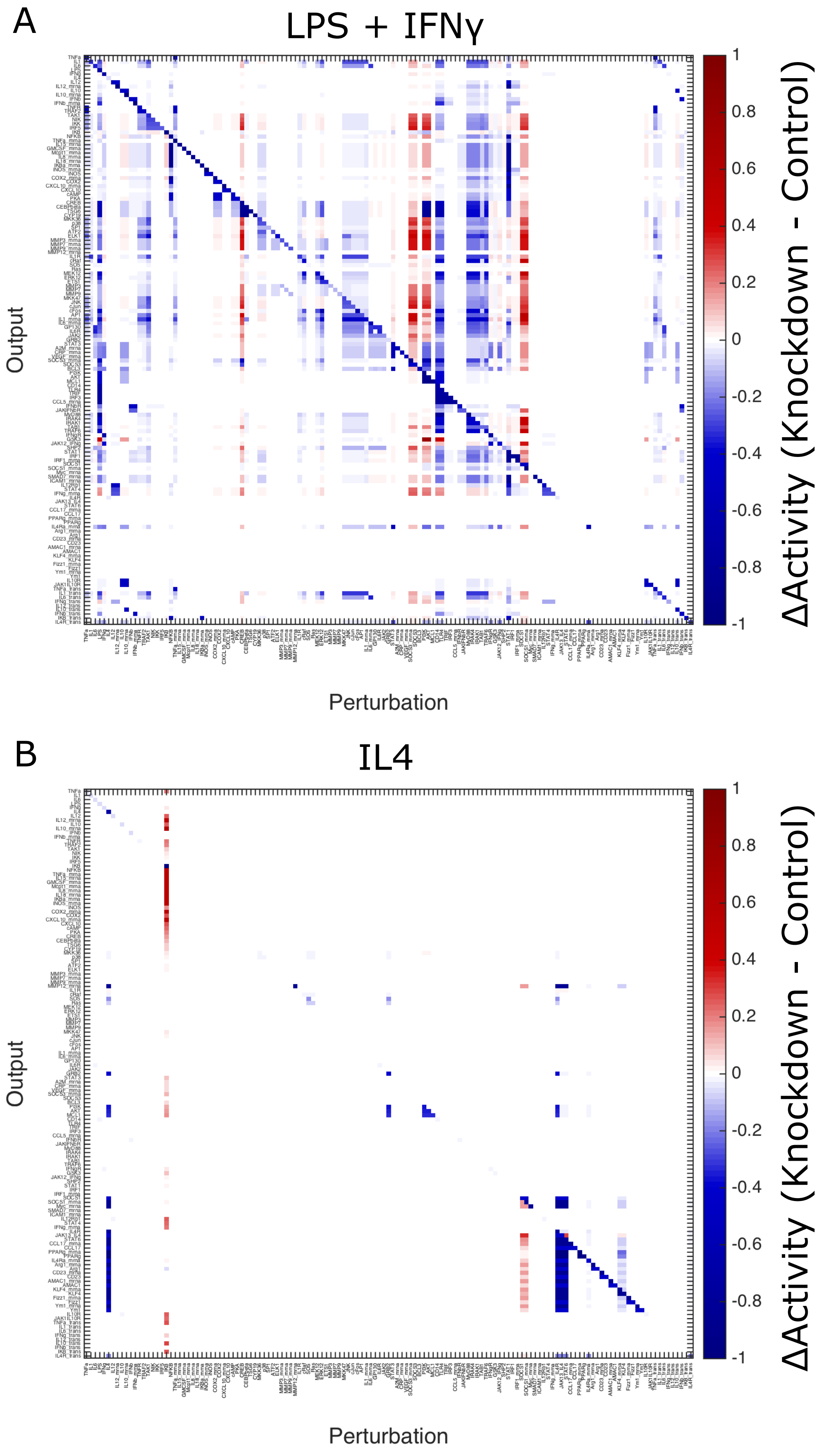
A
LPS + IFNγ
Output
Perturbation
ΔActivity (Knockdown - Control)
1
0.8
0.6
0.4
0.2
0
-0.2
-0.4
-0.6
-0.8
-1
B
IL4
Output
Perturbation
ΔActivity (Knockdown - Control)
1
0.8
0.6
0.4
0.2
0
-0.2
-0.4
-0.6
-0.8
-1

Figure S3

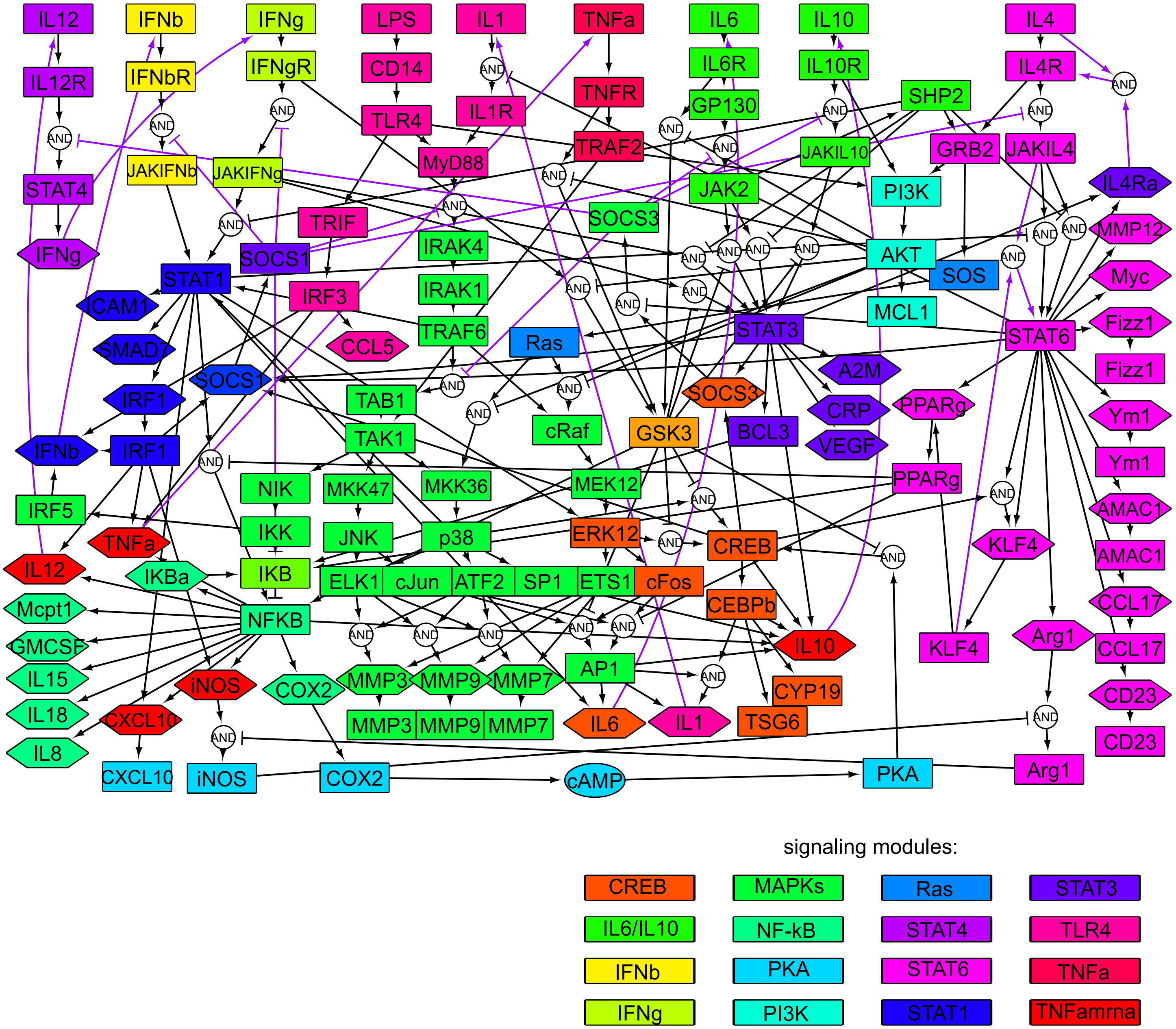

IL12
IFNb
IFNg
LPS
IL1
TNFa
IL6
IL10
IL4
IL12R
IFNbR
IFNgR
CD14
IL1R
TNFR
IL6R
IL10R
IL4R
GP130
SHP2
TLR4
TRAF2
JAKIL10
GRB2
JAKIL4
STAT4
JAKIFNb
JAKIFNg
MyD88
JAK2
PI3K
IL4Ra
TRIF
SOCS3
MMP12
IFNg
IRAK4
AKT
SOS
Myc
STAT1
SOCS1
IRF3
IRAK1
MCL1
ICAM1
TRAF6
STAT3
STAT6
Fizz1
SMAD7
CCL5
Ras
Fizz1
SOCS1
A2M
IRF1
TAB1
SOCS3
CRP
PPARg
Ym1
cRaf
GSK3
BCL3
VEGF
IFNb
IRF1
TAK1
Ym1
NIK
MKK47
MKK36
MEK12
PPARg
IRF5
AMAC1
IKK
JNK
p38
ERK12
CREB
KLF4
TNFa
AMAC1
IL12
IKBa
IKB
ELK1
cJun
ATF2
SP1
ETS1
cFos
CCL17
Mcpt1
NFKB
CEBPb
Arg1
GMCSF
IL10
KLF4
CCL17
MMP3
MMP9
MMP7
AP1
IL15
iNOS
COX2
CYP19
CD23
IL18
CXCL10
MMP3
MMP9
MMP7
IL6
IL1
TSG6
IL8
CD23
CXCL10
iNOS
COX2
cAMP
PKA
Arg1
AND
signaling modules:
CREB
MAPKs
Ras
STAT3
IL6/IL10
NF-kB
STAT4
TLR4
IFNb
PKA
STAT6
TNFa
IFNg
PI3K
STAT1
TNFamrna


Figure S4

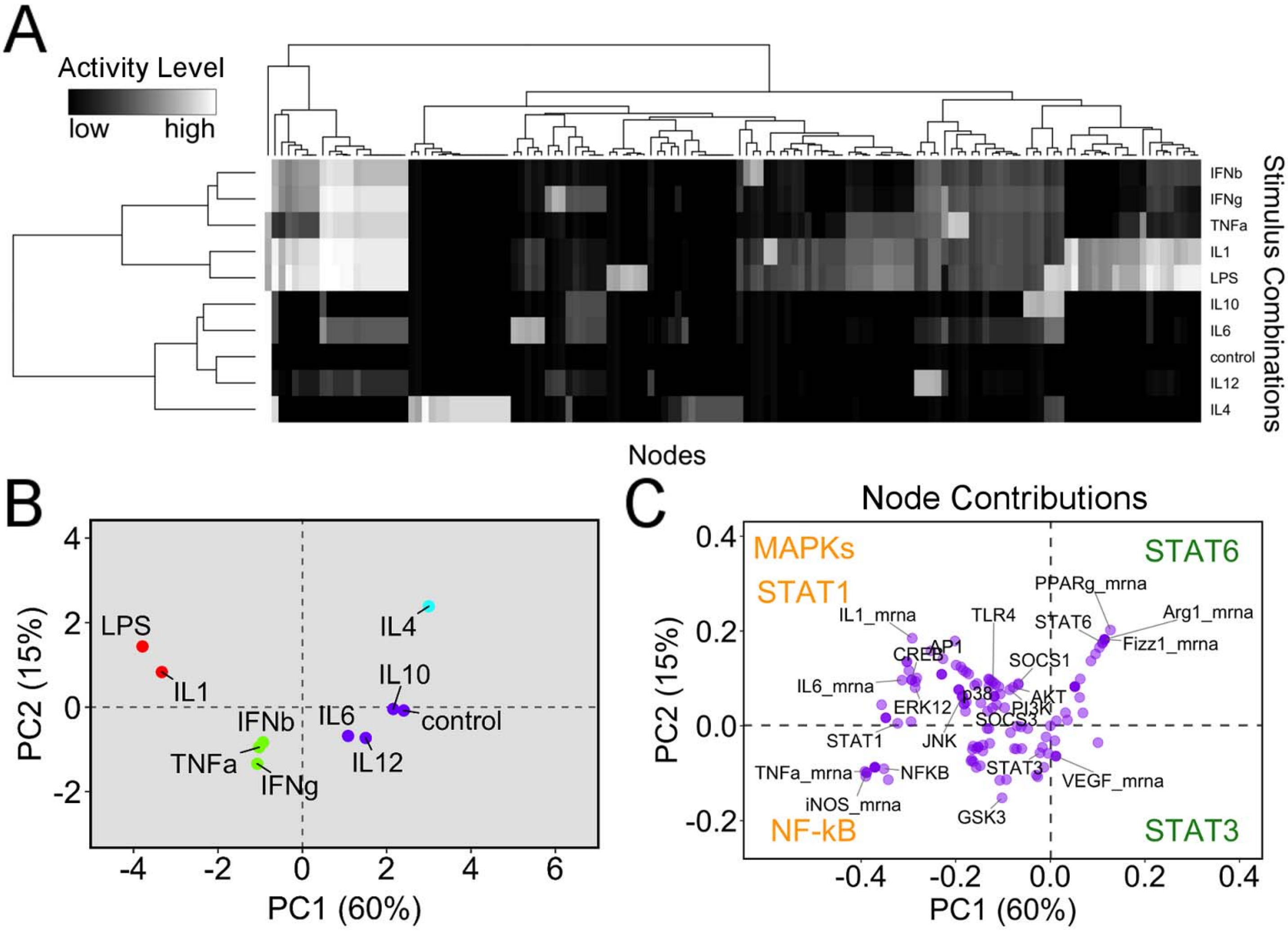

A
Activity Level
low
high
IFNb
IFNg
TNFa
IL1
LPS
IL10
IL6
control
IL12
IL4
Stimulus Combinations
Nodes
B
PC2 (15%)
PC1 (60%)
LPS
IL1
IL4
IL10
IL6
control
IFNb
TNFa
IFNg
IL12
4
2
0
-2
-4
6
C
Node Contributions
PC2 (15%)
PC1 (60%)
MAPKs
STAT1
STAT6
NF-kB
STAT3
PPARg_mrna
Arg1_mrna
STAT6
Fizz1_mrna
IL1_mrna
TLR4
CREB
AP1
SOCS1
IL6_mrna
p38
AKT
ERK12
PI3K
SOCS3
STAT1
JNK
TNFa_mrna
NFKB
STAT3
VEGF_mrna
iNOS_mrna
GSK3
0.4
0.2
0.0
-0.2
-0.4


Figure S5

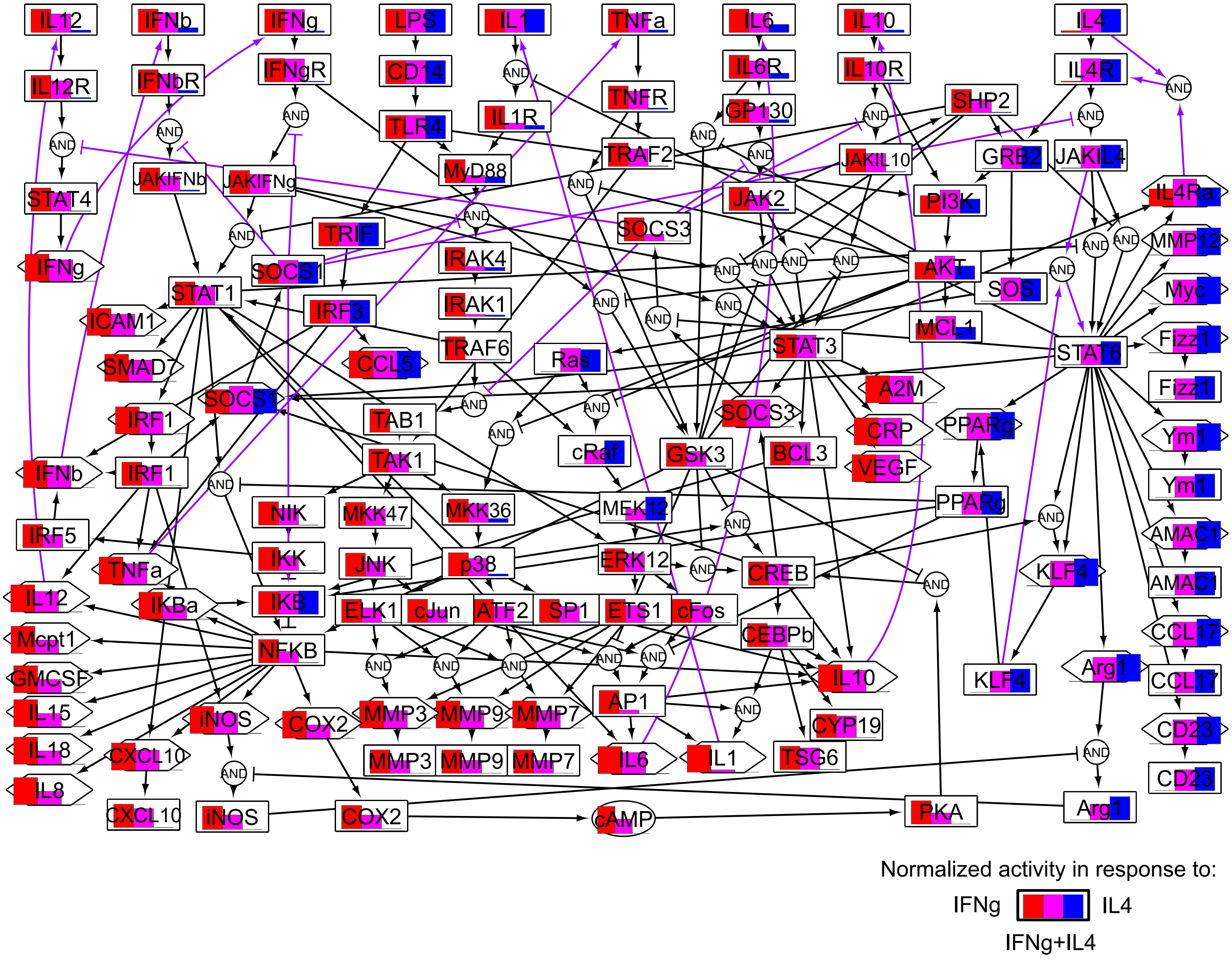
IL12
IFNb
IFNg
LPS
IL1
TNFa
IL6
IL10
IL4
IL12R
IFNbR
IFNgR
CD14
IL1R
TNFR
IL6R
IL10R
IL4R
GP130
SHP2
TLR4
MyD88
TRAF2
JAKIL10
GRB2
JAKIL4
JAKIFNb
JAKIFNg
STAT4
JAK2
PI3K
IL4Ra
TRIF
SOCS3
IRAK4
MMP12
IFNg
SOCS1
AKT
SOS
Myc
STAT1
IRAK1
ICAM1
IRF3
MCL1
Fizz1
TRAF6
STAT3
STAT6
SMAD7
CCL5
Ras
Fizz1
SOCS1
A2M
IRF1
TAB1
SOCS3
CRP
PPARg
Ym1
cRaf
GSK3
BCL3
IFNb
IRF1
TAK1
VEGF
Ym1
MEK12
PPARg
NIK
MKK47
MKK36
IRF5
AMAC1
IKK
JNK
p38
ERK12
CREB
KLF4
AMAC1
TNFa
IL12
IKBa
IKB
ELK1
cJun
ATF2
SP1
ETS1
cFos
CEBPb
CCL17
Mcpt1
NFKB
IL10
Arg1
KLF4
CCL17
GMCSF
AP1
IL15
iNOS
COX2
MMP3
MMP9
MMP7
CYP19
CD23
IL18
CXCL10
MMP3
MMP9
MMP7
IL6
IL1
TSG6
IL8
CD23
CXCL10
iNOS
COX2
cAMP
PKA
Arg1
AND
Normalized activity in response to:
IFNg
IL4
IFNg+IL4

Figure S5

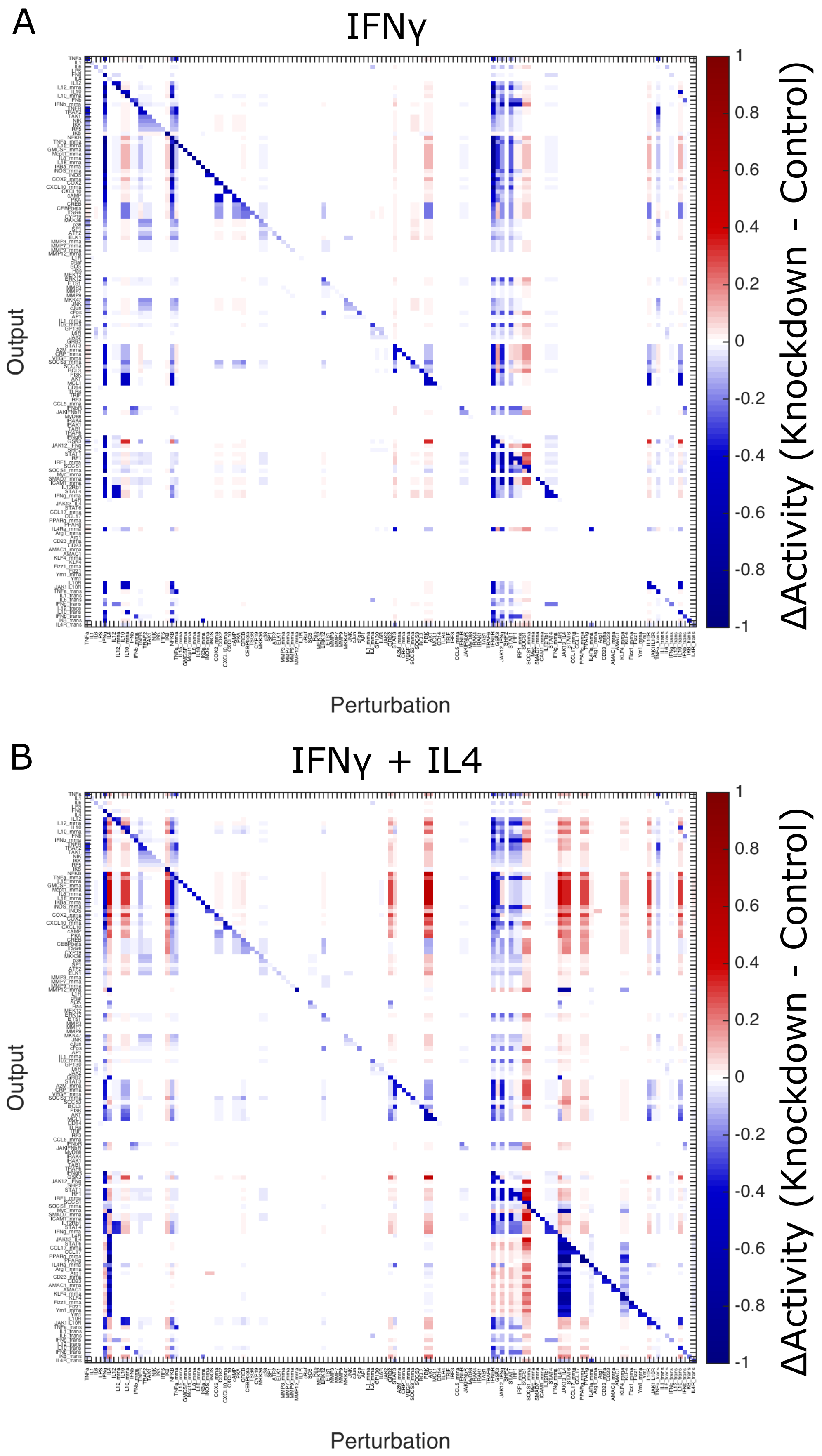
A
IFNγ
Output
Perturbation
ΔActivity (Knockdown - Control)
1
0.8
0.6
0.4
0.2
0
-0.2
-0.4
-0.6
-0.8
-1
B
IFNγ + IL4
Output
Perturbation
ΔActivity (Knockdown - Control)
1
0.8
0.6
0.4
0.2
0
-0.2
-0.4
-0.6
-0.8
-1

Figure S7

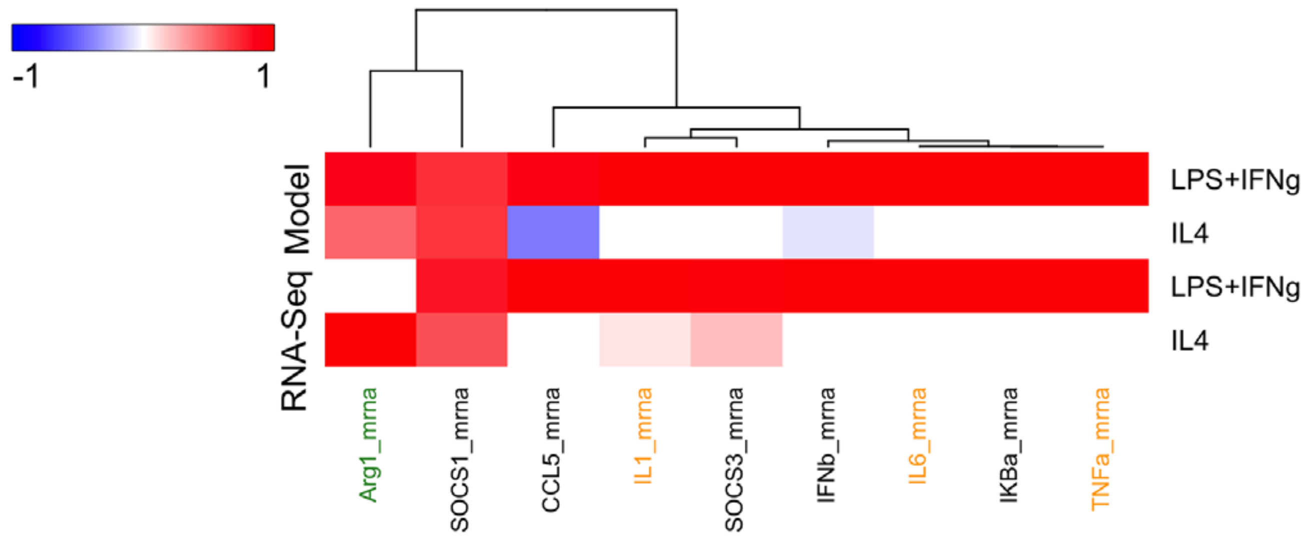

-1
1
RNA-Seq Model
LPS+IFNg
IL4
LPS+IFNg
IL4
Arg1_mrna
SOCS1_mrna
CCL5_mrna
IL1_mrna
SOCS3_mrna
IFNb_mrna
IL6_mrna
IKBa_mrna
TNFa_mrna


Figure S8
